# SegJointGene: joint cell segmentation and spatial gene prioritization by information entropy guided convolutional neural networks

**DOI:** 10.64898/2026.01.08.698474

**Authors:** Haotian Ma, Daifeng Wang

## Abstract

**Motivation:** Spatial sequencing technologies enable the single-cell-level study of molecular organization in tissues. Revealing such spatial patterns relies on accurate cell segmentation. In complex tissues with dense cell packing, segmentation based solely on nuclear staining is insufficient for accurate cell boundary detection. This limitation arises because accurate segmentation necessitates the delineation of cell morphology, which is driven by molecular activities such as cytoskeletal dynamics, cell-cell adhesion, and intercellular signaling. Thus, integrating molecular information, including gene or protein expression, has the potential to improve segmentation, but remains computationally challenging.

**Results:** To address this, we developed SegJointGene, a deep learning framework that jointly performs cell segmentation and spatial gene prioritization by integrating nuclei-based images with spatial gene or protein expression data. SegJointGene designs an information-entropy–guided convolutional neural network together with a computational information discarding score to identify genes that are important for cell-type–specific segmentation. The model iteratively refines gene prioritization and cell boundaries, producing convergent segmentation results along with prioritized spatial genes or proteins across cell types. We applied and benchmarked SegJointGene on various real spatial datasets, including spatial transcriptomics from the mouse hippocampus and distinct regions of the whole mouse brain, as well as spatial proteomics data from human tonsil. Across datasets, SegJointGene outperformed existing methods by 5–20% in accurately assigning molecular signals to cell boundaries. Robustness analyses further demonstrated stable performance across varying gene numbers and imaging resolutions. In addition, the genes prioritized by SegJointGene were enriched for structural, developmental, and synaptic signaling pathways, supporting their relevance to spatial tissue organization.

**Availability:** https://github.com/daifengwanglab/segjointgene

## 1 Introduction

Spatial organization plays a key role in determining cellular function in complex tissues. In the brain cerebral cortex, for example, the laminar position of a neuron affects its connectivity and function within neural circuits. Such spatial organization is fundamentally driven by molecular activities—including cytoskeletal regulation, cell–cell adhesion, and intercellular signaling—that determine cell morphology and define physical boundaries [Eng et al., 2019]. Spatial sequencing technologies have emerged to spatially measure various molecular activities such as gene and protein expression within intact tissue, providing unprecedented access to these spatial patterns at the single-cell level. However, learning such patterns from spatial data depends on accurate cell segmentation, which remains a major technical challenge. Segmentation is particularly difficult in densely packed tissues, where nuclear staining alone is often insufficient to resolve overlapping cells. Moreover, cells with complex morphologies, such as neurons with extended dendritic arbors or glial cells with ramified processes, occupy spatial territories that extend far beyond the nucleus. As a result, nuclear-based segmentation frequently fails to capture true cellular boundaries, leading to ambiguity in assigning cytoplasmic molecular signals.

Although integrating molecular information has great potential for cell segmentation, current computational approaches remain limited. For instance, Cellpose relies primarily on nuclear or membrane staining and does not explicitly leverage spatial gene expression, where most mRNA resides [Stringer et al., 2021]. More recent methods incorporate certain molecular signals directly; for example, JSTA uses gene assignment probabilities to refine cell borders [Littman et al., 2021]. However, such approaches generally treat genes uniformly and do not distinguish morphologically informative genes for various cell types from broadly expressed ones. Similarly, deep learning frameworks such as GeneSegNet [Wang et al., 2023] improve segmentation accuracy by incorporating gene expression features, but they do not explicitly quantify the contribution of individual genes, limiting biological interpretability.

To address these limitations, we developed SegJointGene, a deep learning framework that jointly segments cells and spatially prioritizes genes. Unlike existing approaches that treat molecular signals indiscriminately, SegJointGene incorporates a convolutional neural network guided by a novel computational information discarding (**CID**) score. This interpretable deep learning architecture allows the model to iteratively prioritize morphologically informative genes, identifying the specific molecular markers that drive cell shape while effectively suppressing background noise. We applied SegJointGene to real-world spatial datasets with benchmarks, including mouse brain transcriptomics and human tonsil proteomics, and demonstrated its consistent outperformance across datasets. It improves molecular assignment accuracy in densely packed tissues and reveals cell-type-specific genes associated with structural and signaling functions underlying complex cellular morphologies. SegJointGene is available as an open-source tool at https://github.com/daifengwanglab/segjointgene.

## 2 Methods

As shown in **Fig. 1**, SegJointGene takes two inputs from spatial transcriptomics data: 1) gene density maps **X** derived from mRNA spot coordinates (or proteins from spatial proteomics), and 2) an initial cell segmentation map **A** based on stained cell nuclei and cell-type annotation.

**Figure 1:**
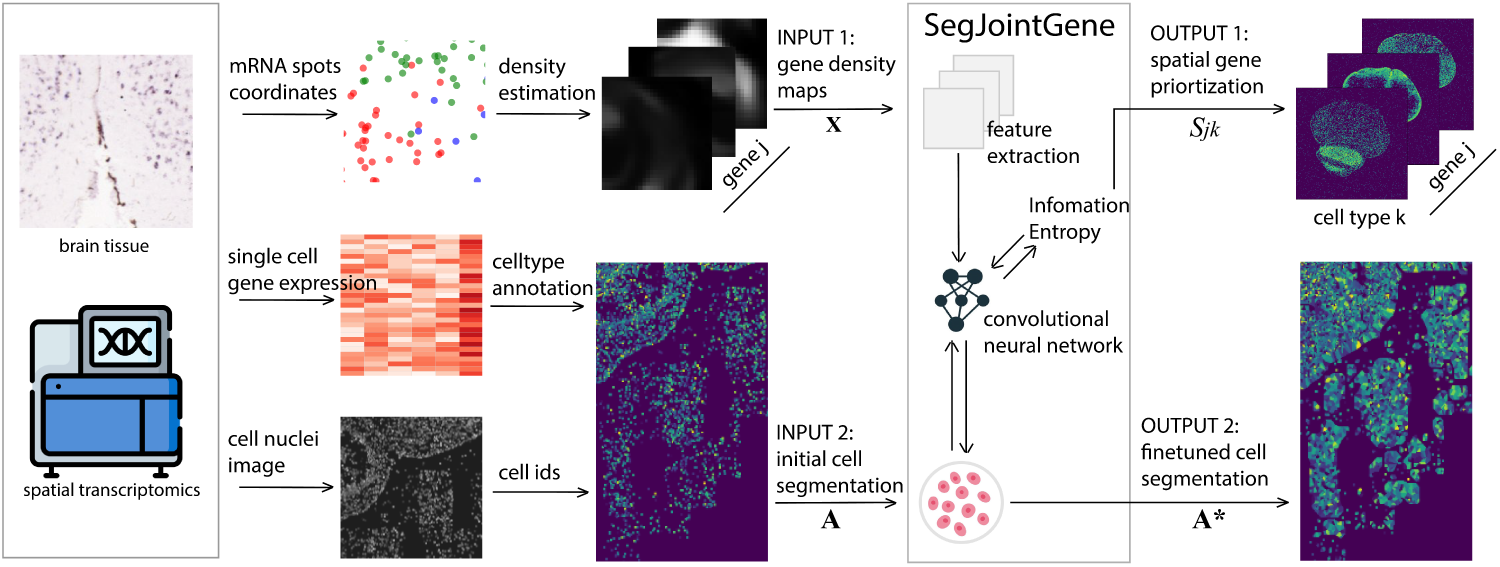
Overview of the SegJointGene framework. The framework integrates spatial transcriptomics to perform joint spatial gene prioritization and cell-segmentation refinement. Input preparation: Spatial transcriptomics data (mRNA spot coordinates, single-cell gene expression, and cell-nuclei images) are processed into two inputs. Input 1: **X** is the stack of gene-density maps derived from mRNA spot coordinates, where each map **X***_j_* represents the spatial distribution of gene *j*. Input 2: **A** is the initial cell-segmentation map derived from cell-nuclei staining and cell-type annotation, assigning a cell-type label *k* to each cell region. SegJointGene: A convolutional neural network (CNN) inputs the gene-density maps **X** and the initial cell-segmentation map **A** as input. The network employs feature extraction and information-entropy calculation to iteratively optimize the segmentation parameters. The framework generates two outputs: Output 1: *S_jk_*, the Gene Importance Score quantifying the spatial relevance of gene *j* for identifying cell type *k*; and Output 2: **A**^∗^, the final cell-segmentation map with refined cell-type assignments *k*.

SegJointGene then iteratively extracts the features using a convolutional neural network (CNN) and updates the cell segmentation map and gene importance scores using information entropy. This process repeats until the segmentation converges. Upon convergence, the framework produces two outputs: 1) a final optimized segmentation map **A**^∗^, and 2) spatial gene importance scores for cell type-level segmentation, i.e. *S_jk_*for gene *j* and cell type *k*.

We define the variables of our algorithm as follows. Let **A** ∈ ℤ*^w^*^×*l*^ be the cell segmentation map for an image of width *w* and length *l*, for a total of *N* = *w* × *l* pixels. Each entry *A_i_* ∈ {1*,…, K*} contains the cell-type label for the pixel *i*. Let **X** ∈ ℝ*^w^*^×*l*×*M*^ be the stack of *M* gene density maps, where **X***_j_* represents the density map for gene *j* (**Supplementary Note S1**).

As detailed in **Algorithm 1**, the core procedure alternates between model optimization and segmentation refinement. In each iteration, we first train a CNN parameterized by ***θ*** to predict cell types using the current map **A** as the ground truth. Next, we compute pixel-specific importance scores *S_jki_* using CID (Section 2.3.1) and aggregate them into a metric *I_ik_*to quantify the relevance of local features. Finally, the segmentation map **A** is updated by reassigning border pixels where both the confidence of the model (*p_ik_*) and the aggregate importance (*I_ik_*) exceed their respective thresholds. The loop ends when the update rate falls below *ɛ*.

**Algorithm 1.**
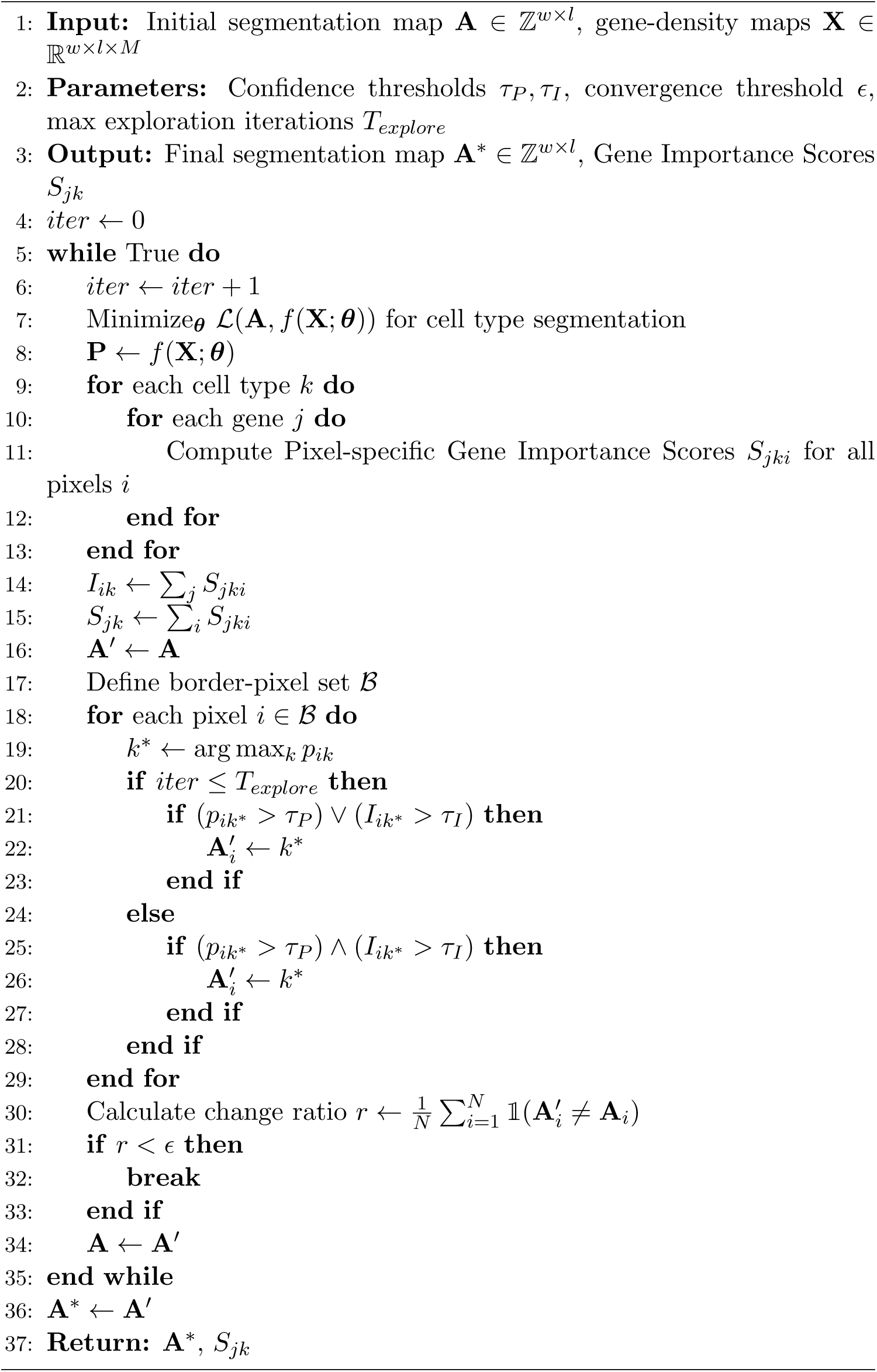
SegJointGene algorithm.

### 2.1 Step 1: Training the segmentation model

Cell segmentation here is formulated as a pixel-wise classification problem. The goal is to assign a cell type label *k* ∈ {1, 2*,…, K*} to every pixel *i* in an input image. Here, *K* is the total number of cell types, including the background.

For this task, we employ a U-Net architecture as our convolutional neural network backbone *f* (·; ***θ***). The U-Net is exceptionally well-suited for biomedical image segmentation because of its design. It consists of an encoder that captures contextual information from input gene maps and a decoder that enables precise localization (**Supplementary Note S2**). Skip connections between the encoder and decoder paths are a key feature, allowing the network to combine high-level feature information with fine-grained spatial details, which is critical for accurately delineating cell boundaries.

The input to the U-Net is a tensor of size *w* × *l* × *M*, representing the image dimensions and the *M* gene density maps. The network outputs a probability tensor of size *w* × *l* × *K*. The final layer of the network is a soft-max function, which computes the probability *p_ik_* that the pixel *i* belongs to the cell type *k*. We train the network parameters ***θ*** by minimizing the discrepancy between the prediction of the model and the current segmentation map **A**, serving as the ground truth for this iteration:

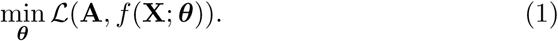

Specifically, the loss function 𝒧 is defined as the Categorical Cross-Entropy loss summed over all pixels:

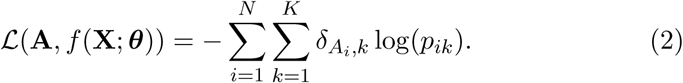

In this formula, *δ_Ai,k_* is the Kronecker delta; it is 1 if the true integer label of the pixel *i* is the cell type *k* and 0 otherwise. Functionally, this term acts as a selector that filters out all incorrect classes from the summation, retaining only the log-probability of the true cell type. The minimization of this loss updates the model parameters ***θ***, which are utilized in Step 2 to calculate gene importance scores.

### 2.2 Step 2: Updating segmentation by flipping border pixels

While the initial U-Net training provides a baseline segmentation, a standard end-to-end approach is insufficient for this task due to the fundamental lack of a ground truth signal in the extranuclear space. Our primary goal is to leverage spatial patterns in gene expression to accurately infer cell boundaries beyond the initial nuclei. To achieve this, we introduce an iterative, model-guided refinement process that focuses on re-assigning the labels of border pixels, which we define as the set of all pixels, B, adjacent to a currently labeled cell region. We explore several strategies for this refinement.

#### 2.2.1 Refinement based on predictive probability

The most direct method for updating border pixels is to use the raw predictive probabilities from the model’s softmax output. For each border pixel *i* ∈ B, the trained model *f* (**X**; ***θ***) generates a probability vector **p***_i_* = [*p_i_*_1_*, p_i_*_2_*,…, p_iK_*]^⊤^ across all *K* cell types. Let *k*^∗^ = arg max*_k_ p_ik_* be the cell type with the maximum predicted probability. To prevent low-confidence guesses from corrupting segmentation, we introduce a confidence threshold, *τ_P_*. The pixel’s label is updated only if the maximal probability exceeds this threshold:

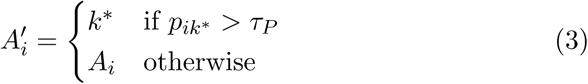

Although straightforward, this approach can be unreliable in ambiguous regions where the model may output high probabilities that are not strongly supported by the underlying gene expression evidence.

#### 2.2.2 Enhancing cell segmentation by aggregated pixel importance score

To create a more robust update rule, we use a feature attribution method based on the Deep InfoDiscard framework [Ma et al., 2022], as detailed in Section 2.3. This method calculates an Aggregated Pixel Importance Score, *I_ik_*, which directly quantifies how much the local gene expression patterns contributed to the model’s decision to classify pixel *i* as cell type *k*. We hypothesize that this score is a more faithful measure for a given classification. In this strategy, the potential update is still determined by the most probable class *k*^∗^ as defined above, but the assignment is contingent on its associated Aggregated Pixel Importance Score passing an importance threshold *τ_I_*:

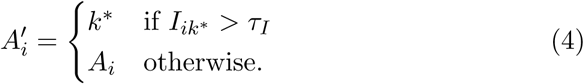

This importance-driven approach ensures that the changes are biologically significant. However, relying solely on importance can be overly conservative, potentially missing valid boundary expansions that are confidently predicted by the model. Therefore, to achieve an optimal balance between confident exploration and robust refinement, we developed a combined two-phase strategy.

#### 2.2.3 Two-phase strategy for updating cell boundaries

To balance boundary exploration in the early stages with high-precision refinement in the later stages, we implemented a mixed strategy that evolves with the iterative process.

- **Exploration phase:** In initial iterations, the goal is to allow the cell boundaries to expand from the confident nuclei. We use a more lenient update rule where a pixel’s label is flipped if it satisfies a disjunctive condition: the update is triggered if either the maximal predictive probability exceeds *τ_P_* or the maximal Aggregated Pixel Importance Score exceeds *τ_I_*.
- **Convergence phase:** As the segmentation matures, the priority shifts to ensuring high fidelity and stability. To achieve this, the update rule becomes more stringent. The label of a pixel is reassigned only if both criteria are met simultaneously: the predictive probability must exceed *τ_P_*and the aggregate Pixel Importance Score must exceed *τ_I_*. This dual confirmation requirement prevents noisy fluctuations at the boundaries and ensures that only the most confident and biologically-supported changes are made, leading to a stable and high-accuracy convergence.

Critically, this updated map **A**^′^ serves as the new ground truth for the subsequent iteration of Step 1. By replacing the target **A** in Equation 2, the optimization objective dynamically changes, requiring the model to learn features from the newly discovered cytoplasmic regions rather than just the initial nuclei.

### 2.3 Spatial gene prioritization via information entropy theory

To obtain a robust importance score for our refinement step, we extended the Deep InfoDiscard framework to spatial omics data. Traditional attribution methods often cannot be fairly compared across different network layers or architectures. The Deep InfoDiscard method addresses this by introducing a theoretically grounded metric based on information theory to quantify how much information about a given input pixel is discarded or ignored by the network during computation.

#### 2.3.1 Computational information discarding

The core metric we used is CID, which measures the network’s invariance to a specific input gene map when identifying a particular cell type *k* (**Fig. 2**). Conceptually, it seeks the maximum amount of noise that can be added to the density map of a single gene (e.g. **X***_j_*) while maintaining the prediction of network output. This total CID score decomposes into pixel-wise entropies *H*(*σ_jki_*), which we visualized to reveal local spatial relevance.

**Figure 2:**
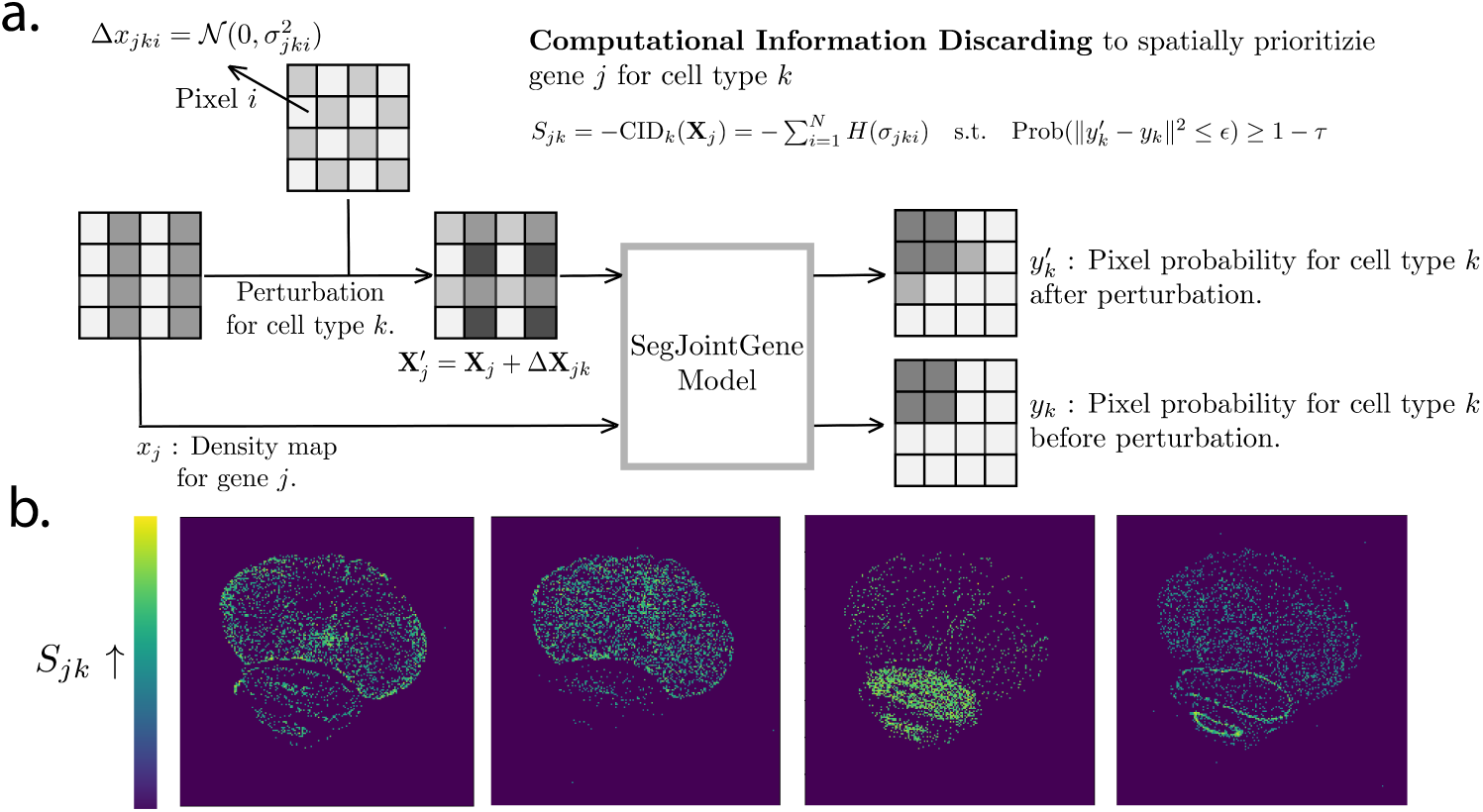
Information discarding for spatial gene–cell-type prioritization. **a**, Illustration of Computational Information Discarding (CID), a method adapted to quantify the *invariance* of a model’s cell-type prediction to spatial gene expression data. The schematic shows how CID quantifies the maximum noise entropy, CID*_k_*(**X***_j_*), that can be added to an input 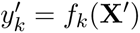 while keeping the model’s output for a specific cell type 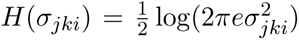 stable and close to the original *y_k_*. Here, **X***_j_* is the density map for gene *j*, and Δ*x_jk,i_* is the perturbation applied to pixel *i*. A high CID*_k_*(**X***_j_*) score signifies that the model’s prediction for cell type *k* is highly invariant to (i.e., irrelevant to) the expression of gene *j*. **b**, The bottom heatmaps visualize the spatial distribution of pixel specific gene importance score *S_jk_*, computed by CID. These maps illustrate the local irrelevance of a gene’s expression to a cell type’s classification. High-CID (e.g., yellow) regions are irrelevant, whereas low-CID (e.g., purple/blue) regions are relevant.

Let **X** be the complete input stack of all *M* gene-density maps, and let **X***_j_* be the specific density map for gene *j*. Let *y_k_* = *f_k_*(**X**) be the original output probability for cell type *k*. To calculate CID for the gene *j*, we perturb only its map by adding Gaussian noise to pixels, creating a perturbed map 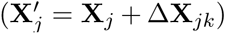, where Δ**X***_jk_* ∼ 𝒩 (0, **Σ***_jk_*). Here, **Σ***_jk_* is a diagonal covariance matrix with variances 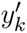. The vector ***σ****_jk_* = [*σ_jk_*_1_*,…, σ_jkN_*] represents the standard deviations in pixels. The entire input stack temporarily becomes 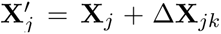.

The objective is to maximize the entropy of this noise, subject to the constraint that the new output 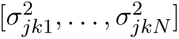 remains close to the original *y_k_*. We formally establish that the Gene Importance Score corresponds to the negative of the total Information Discarding *S_jk_* = −*CID_k_*(**X***_j_*). The CID metric itself is defined as follows:

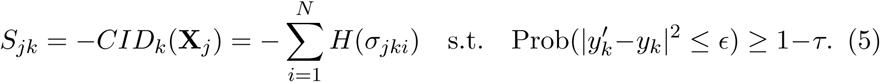

Here, the pixel-wise entropy is 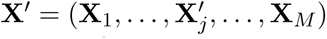, where *σ_jki_* is the optimal standard deviation for the pixel *i* of gene *j* when constrained by cell type *k*. A high *H*(*σ_jki_*) indicates that the prediction of the model is highly invariant with the expression of gene *j* in the pixel *i*. In contrast, a low entropy indicates high local relevance.

To solve this constrained optimization problem, we formulate a tractable loss function that is minimized through gradient descent to find the optimal standard deviations ***σ****_jk_*for gene *j* and cell type *k*:

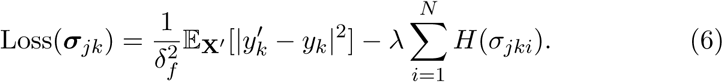

The first term acts as an error penalty, measuring how much the perturbed output *f* (**X**^′^) deviates from the original output for cell type *k*. The second term works to maximize the input entropy of the gene *j*.

#### 2.3.2 Quantifying spatial gene importance via CID

A large *σ_jki_* means that the network can tolerate significant noise in that pixel without altering its internal representation of a cell type; therefore, the information in that pixel is discarded. In contrast, a small *σ_jki_* means that the network is highly sensitive to the value in that pixel, implying that its information is critical to the prediction.

This works because, unlike raw softmax probabilities which only reflect the final guess of the model, the CID score directly probes the internal mechanism of the model and its dependence on specific input features. In our work, the Aggregated Pixel Importance Score *I_ik_* is derived directly from these pixel-wise entropies: *I_ik_* =Σ*_j_ S_jki_* with *S_jki_* = −*H*(*σ_jki_*). Pixels where genes are highly informative for cell type *k* have low pixel-wise entropies *H*(*σ_jki_*), which, when summed and negated, yield high positive importance scores used in refinement.

### 2.4 Evaluation

To quantitatively assess the performance of our segmentation, we adopted the cell calling score, a metric previously used by GeneSegNet that measures the ability of a method to correctly assign molecules to the interior of a predicted cell. Let *G_in_*be the set of molecules located within a predicted cell region and *G_out_* be the set of molecules located in the immediate exterior of a cell, specifically within a distance threshold (*δ*) of the predicted boundary. The cell calling score is calculated as:

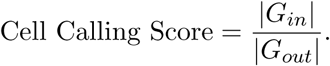

A higher score indicates a stronger ability to recruit molecules within predicted cell boundaries while excluding those just outside, signifying a more precise segmentation. For our analysis, we used distance thresholds of 3, 5, and 7 pixels. This metric is particularly effective for crowded tissue regions where RNAs from neighboring cells can form a continuum, making simple distance-based assignment difficult.

### 2.5 Datasets and data processing

To evaluate the performance and robustness of SegJointGene, we applied our method to three distinct datasets representing different biological contexts. First, we benchmarked SegJointGene on a publicly available data set of the mouse hippocampal CA1 region, previously used to evaluate GeneSegNet [Wang et al., 2023]. This dataset, generated via in situ RNA detection, is characterized by sparse RNA reads, providing a challenging scenario to test the algorithm’s sensitivity in data-scarce conditions.

Second, we applied our method to the widely recognized the Whole Mouse Brain dataset from the Allen Institute for Brain Science [Yao et al., 2023]. This data set features a highly complex and heterogeneous tissue environment with a dense arrangement of diverse cell types, allowing us to assess the performance and scalability of SegJointGene in a large-scale and intricate sample.

Finally, to demonstrate the adaptability of the SegJointGene framework beyond transcriptomics, we analyzed a human tonsil spatial proteomics dataset containing Imaging Mass Cytometry (IMC) data [Lee et al., 2025]. In this application, we replaced the gene-density maps with protein-density maps, testing the algorithm’s ability to leverage alternative types of spatially resolved molecular data for accurate cell segmentation.

For all datasets, input expression data were standardized using a z-score transformation. Since these datasets were pre-curated, no additional quality control or preprocessing was required. If a dataset consisted of a single tissue section, we randomly split the image grids from that section into 70% training, 10% validation and 20% testing sets; otherwise, data from different tissue sections were divided into training, validation, and testing sets according to the same ratio.

### 2.6 Gene set enrichment analysis

To interpret the biological significance of the genes prioritized by our method, we performed pathway and process enrichment analysis using the Metascape web server [Zhou et al., 2019]. For the analysis, the complete set of all input genes was used as the statistical background. We then computed functional enrichments for the list of prioritized genes identified by SegJointGene. This ensures that the identified pathways are specifically enriched within the subset of genes that our model found most informative for the segmentation task, rather than reflecting general properties of the full gene list. However, due to the limited number of genes in the mouse hippocampus dataset (84 genes), using them as background was statistically impractical. Instead, we utilized the entire mouse genome as the background to ensure more robust enrichment results.

## 3 Results

### 3.1 Synaptic and neurotransmission genes for spatially enhancing cell segmentation in the mouse hippocampus

First, we benchmarked and evaluated SegJointGene and other methods using the cell calling score for accurate mRNA spot assignment using a spatial transcriptomics dataset in the mouse hippocampus [Wang et al., 2023] (**Fig. 3a, Methods**). SegJointGene demonstrated superior performance across all pixel distances evaluated (3, 5, and 7 pixels). For example, according to the 3-pixel criterion, SegJointGene outperformed GeneSegNet by 3.99, JSTA by 24.3 and Cellpose by 36.99. The cell calling score converges through iterations (**Supplementary Fig. S1**).

**Figure 3:**
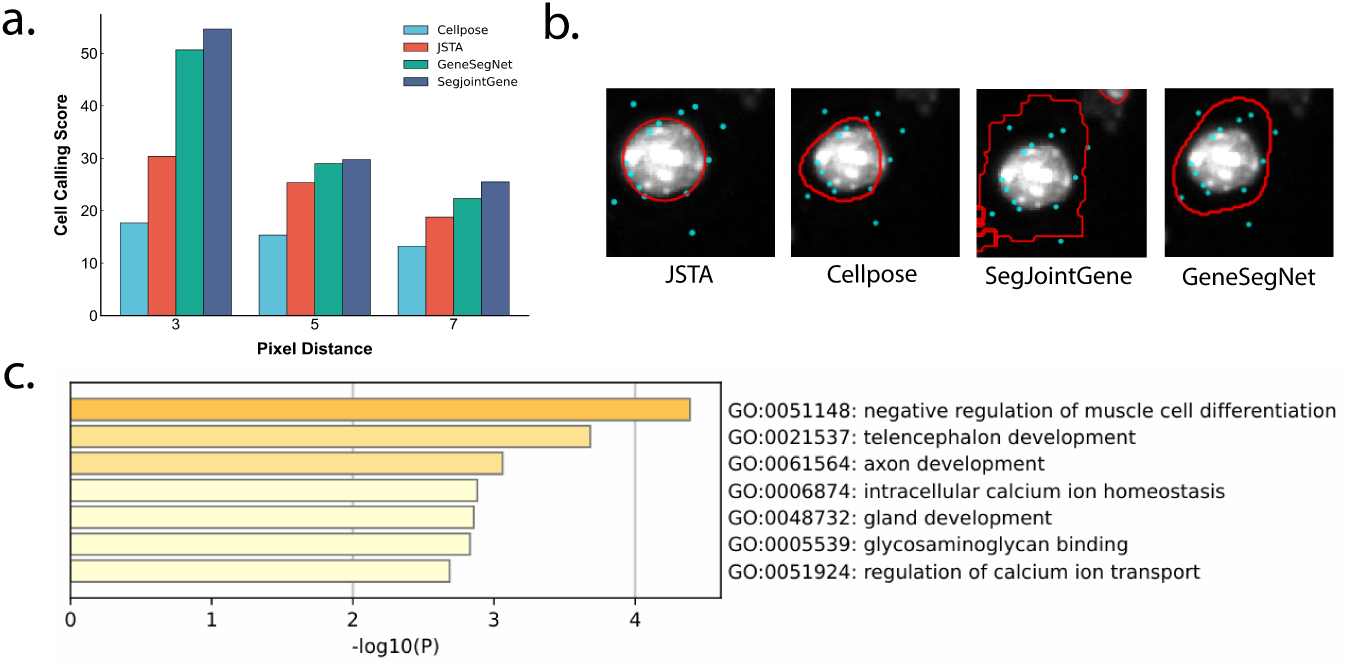
Benchmarking, cell border visualization, and gene enrichment analysis of SegJointGene in the mouse hippocampus. **(a)** Quantitative comparison of SegJointGene with existing segmentation approaches using the cell calling score across multiple pixel-distance thresholds (3, 5, and 7 pixels). Higher scores indicate more accurate mRNA-to-cell assignment. **(b)** Visualization of cell boundaries from the CA1 region of mouse hippocumpus. Cyan: mRNA spots. Red: cell border predicted by methods. Gray: cell nuclei. **(c)** Gene set enrichment analysis of the top 30% prioritized genes (25 out of 84) identified by SegJointGene.

We also visualized cell borders for select cells with high cell calling scores to qualitatively support these quantitative evaluations (**Fig. 3b**). The boundaries generated by SegJointGene (red) aligned closely with the clusters of mRNA points (cyan), whereas the reference methods incorrectly truncated these cell bodies or defined their boundaries irregularly. By integrating gene density maps with cell nucleus boundaries, SegJointGene provided highly accurate delineation of both large pyramidal neurons with extended cytoplasm and complex interneurons with irregular shapes. This precision helped ensure an accurate assignment of mRNA spots, preventing their misallocation to neighboring cells or the extracellular space.

Moreover, enrichment analysis of the genes prioritized by SegJointGene revealed a significant association with neurodevelopmental and calcium-regulated neuronal processes (*p <* 10^−3^) (**Fig. 3c**, Supplementary Data 1). These genes were not selected only based on expression magnitude or cell-type specificity, but were prioritized by their contribution to segmentation of cellular boundaries. Consequently, the enriched terms were dominated by pathways related to telencephalon and axon development, as well as regulation of calcium ion transport and intracellular calcium homeostasis—processes that are intrinsically spatially structured and closely coupled with neuronal morphology, differentiation, and functional compartmentalization in the hippocampus [Diethorn and Gould, 2023]. Such processes are known to form the basis for adult hippocampal neurogenesis [Goņcalves et al., 2016], circuit integration, and activity-dependent transcriptional regulation [Greer and Greenberg, 2008], all of which rely on precise cellular organization and boundary definition during neural development.

### 3.2 Cross-region spatial gene prioritization for cell segmentation in the Whole Mouse Brain dataset

Next, to evaluate robustness across various brain regions, we applied SegJointGene to a whole mouse brain dataset (**Fig. 4a**). Consistent with the hippocampus results, our method maintained superior performance. Specifically, under the stringent 3-pixel threshold, SegJointGene surpassed the next-best performer, GeneSegNet, by approximately 1.82 (SegJointGene: 32.87 vs. GeneSegNet: 31.05). This highlighted the model’s capacity to handle complex, brain-wide anatomical structures. Moreover, SegJointGene demonstrated robust outperformances across varying input gene numbers (**Supplementary Fig. S2**).

**Figure 4:**
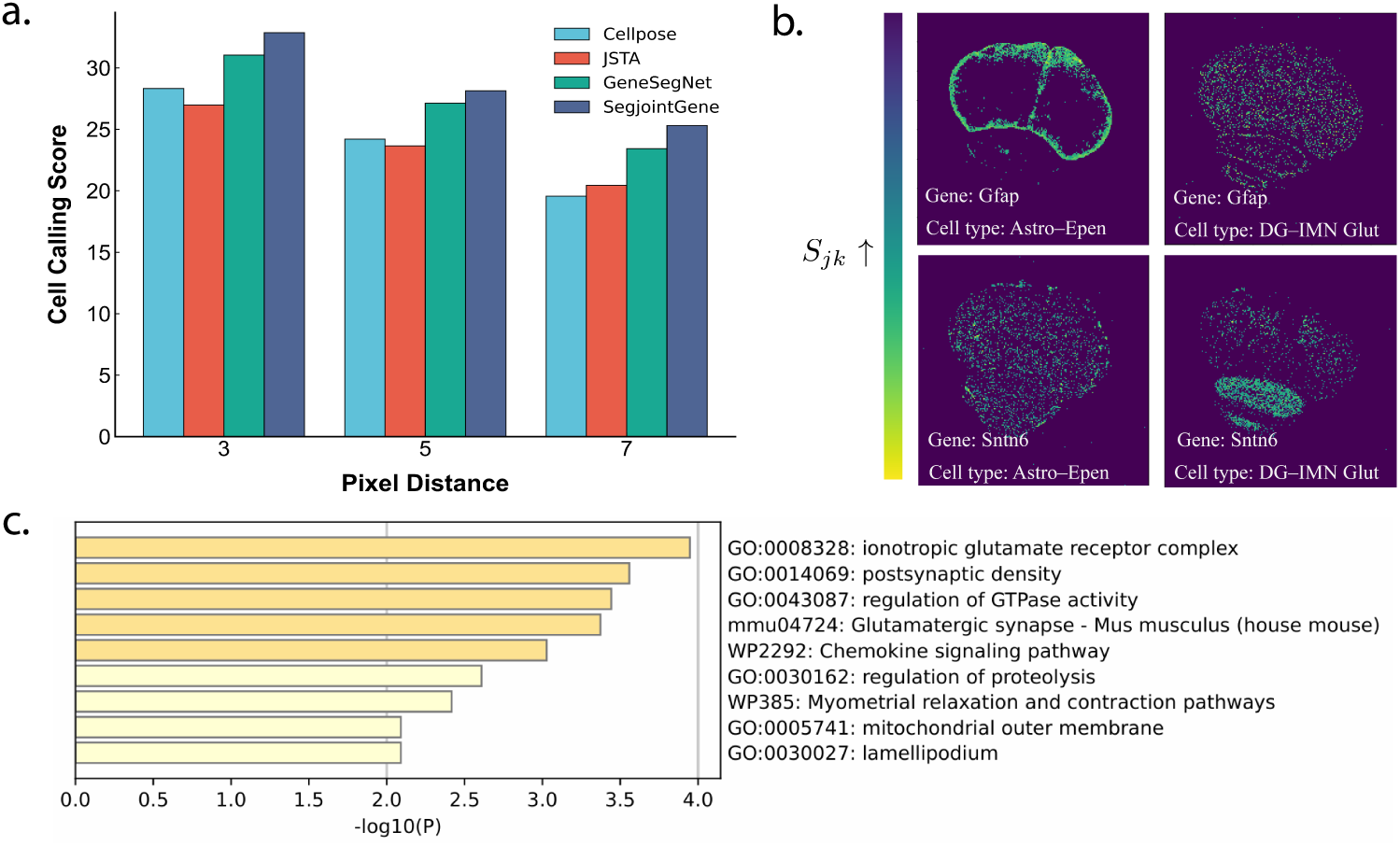
Benchmarking, importance score visualization, and gene enrichment analysis of SegJointGene on the Whole Mouse Brain dataset. **(a)** Benchmarking of cell calling scores across different methods under varying pixel distance thresholds (3, 5, and 7 pixels). **(b)** Spatial visualization of Gene Importance Scores in a coronal mouse brain section. The maps reveal how the model identifies distinct anatomical structures based on specific gene-cell type pairs: *Gfap* (for cell type Astro-Epen) highlights the ventricular and peri-ventricular boundaries, while *Syt6* (for cell type DG-IMN Glut) specifically localizes to the hippocampus and dentate gyrus in the ventral region. **(c)** Gene enrichment analysis of the top 10% genes (30 out of 300) prioritized by SegJointGene in the mouse prefrontal cortex.

Local CID maps illustrated how SegJointGene captured gene-cell-type specificity across distinct regions (**Fig. 4b**). Gfap showed clear spatial relevance around the ventricular and peri-ventricular structures, consistent with the structural roles of the glial and ependymal populations in maintaining the organization of the epithelium [van Roy and Berx, 2008]. In contrast, Syt6 exhibited strong relevance within the hippocampus and the dentate gyrus enriched for glutamatergic granule neurons, which are characterized by high synaptic activity and expression of synaptic proteins [Tremblay et al., 2016, Malenka and Bear, 2004]. These spatially coherent patterns demonstrated that SegJointGene recovered biologically grounded gene-cell-type relationships that aligned with known neuroanatomy.

To confirm that SegJointGene’s prioritization yields biologically meaningful results in a different and highly complex brain region, we performed a gene set enrichment analysis on the top 10% of genes prioritized from the mouse prefrontal cortex (**Fig. 4c**, **Supplementary Data 2**). The analysis revealed a significant enrichment for the pathways critical to neuronal connectivity and signal transduction (*p <* 10^−3^). The key enriched terms highlighted components integral to the synaptic interface, such as the ionotropic glutamate receptor complex and the postsynaptic density, which physically define the communicative boundaries between neurons [Sheng and Kim, 2011, Sheng and Hoogenraad, 2007]. Furthermore, the enrichment of lamellipodium indicates the model’s ability to capture dynamic morphological edges driven by cytoskeletal extensions [Pollard and Borisy, 2003]. This finding underscores SegJointGene’s capacity to identify a functionally coherent set of surface-associated markers, effectively delineating the complex physical and functional borders of cells in the prefrontal cortex.

### 3.3 Prioritizing cell segmentation marker proteins over expression-based markers in the human tonsil

To evaluate the generalizability of SegJointGene across different molecular modalities, we assessed its performance on spatial proteomics data acquired via Imaging Mass Cytometry (IMC). Benchmarking on these data demonstrated that SegJointGene significantly outperformed other methods (**Fig. 5a**). In particular, at a 5-pixel distance, it achieved a score of 13.46, substantially surpassing JSTA (5.24), Cellpose (7.09) and GeneSegNet (10.99), confirming its effectiveness in protein-rich environments.

**Figure 5:**
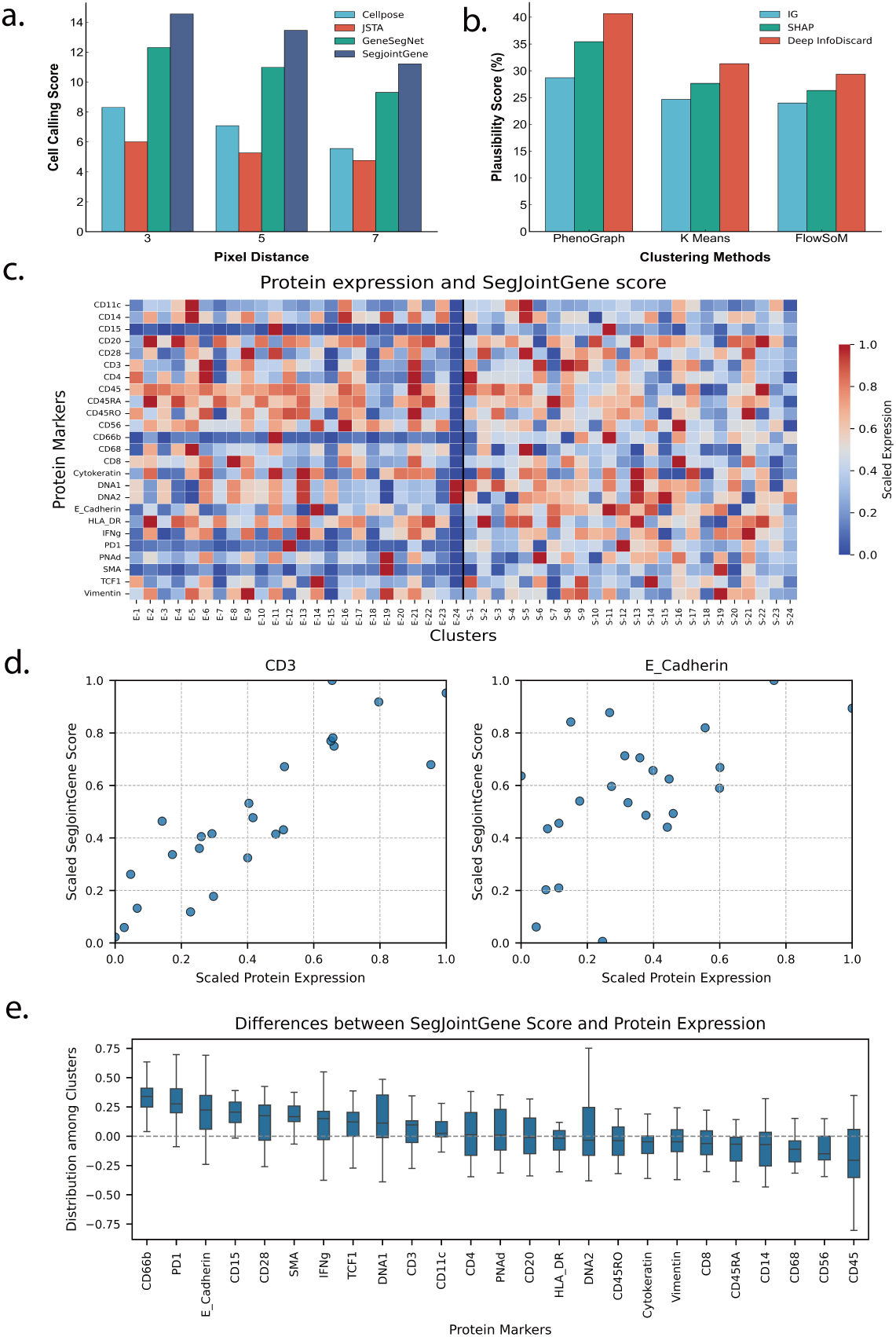
Benchmarking and importance score evaluation of SegJointGene on a spatial proteomics dataset from human tonsil. **(a)** Benchmarking of cell calling scores across different methods. **(b)** Benchmarking of the biological plausibility scores for protein patterns identified by Deep InfoDiscard against Intergrated Gradient (IG) and SHAP. **(c)** Heatmap showing SegJointGene scores and protein expression across clustered cell types. **(d)** Scatter plots for representative markers, showing the positive correlation between SegJointGene scores and protein expression. **(e)** Bar chart of the average difference between SegJointGene scores and protein expression for each protein.

In addition, we measured the biological meanings of the protein features prioritized by SegJointGene. We used the plausibility score, a metric designed to quantify the significance of coexpression patterns among proteins. To show our strength, we compared the patterns derived from SegJointGene’s feature importance scores against other interpretation methods (**Fig. 5b**). When combined with standard clustering algorithms like PhenoGraph, SegJointGene’s prioritized proteins achieved a plausibility score of 40.67%, better than the scores from other interpretation techniques such as IG (15.66%) [Sundararajan et al., 2017] and SHAP (35.43%) [Lundberg and Lee, 2017]. This indicated that SegJointGene could do better protein prioritization for clustering, and thus bring more plausible biological relationships among proteins.

Finally, to visually assess the biological relevance of the prioritized proteins, we compared the raw scaled expression of each marker with its corresponding SegJointGene importance score in 24 distinct cell groups (**Fig. 5c**). The dual heat map revealed a strong concordance between protein expression patterns and SegJointGene scores, indicating that the model successfully identified biologically significant markers.

Proteins that are clearly useful for defining the cell structure consistently received high importance scores (**Supplementary Data 3**). For example, DNA1 and DNA2, which mark the cell nucleus, and E-Cadherin, a canonical cell-cell adhesion molecule essential for maintaining epithelial tissue structure [van Roy and Berx, 2008], were assigned uniformly high scores. Similarly, CD45, a pan-leukocyte marker critical for defining the boundaries of immune cells within the tissue microenvironment [Hermiston et al., 2002], was also correctly identified as highly important. Furthermore, key cell lineage markers that define the architecture of distinct cell neighborhoods—such as CD3 as a definitive marker for the T cell lineage [Smith-Garvin et al., 2009], CD20 for B-cells [Coss and El-Masry, 2024], and Cytokeratin to provide structural identity for epithelial cells [Moll et al., 2008]—were assigned high scores. This demonstrated that SegJointGene effectively learned to prioritize the proteins that are crucial for defining a cell’s identity and its physical boundaries.

We plotted the SegJointGene importance score against the scaled protein expression for each of the 24 cell clusters (**Fig. 5d**). For the T-cell lineage marker CD3, the resulting graph revealed a strong positive correlation, with data points clustering tightly around the identity line. This validated the model’s fundamental ability to accurately track the expression of key cell-type-defining markers. More revealing was the analysis for E-Cadherin, a canonical cell-cell adhesion molecule. Although also showing a positive trend, many clusters for E-Cadherin lay distinctly above the identity line, indicating that the model systematically assigned a higher importance score than what its expression level alone would suggest. SegJointGene not only mirrored expression intensity, but actively identified and amplified the significance of proteins that were functionally critical to defining physical cell boundaries, thereby discovering their higher importance in the segmentation task.

To systematically assess this phenomenon across all markers, we computed the average difference between the SegJointGene score and protein expression for each protein (**Fig. 5e**). This analysis uncovered the model’s learned prioritization scheme. Proteins such as E-Cadherin, PD1, and CD66b showed a significant positive difference value, indicating that the model weighted markers were crucial for cell adhesion and intercellular interactions—features essential for delineating cell borders. In contrast, markers such as the pan-leukocyte protein CD45 exhibited a negative difference value. Despite its high and widespread expression, its utility in distinguishing between adjacent subtypes of densely packed immune cells was limited. The model assigned it a lower relative importance, prioritizing markers with greater distinguishing power over those with signal intensity. Ultimately, it demonstrated that SegJointGene implemented a non-linear, biologically-informed weighting logic, learning to prioritize proteins based on their functional context and utility in defining the cellular architecture of the tissue.

## 4 Discussion

In this work, we introduced SegJointGene, a deep learning framework that significantly advances cell segmentation for spatial omics data. The primary innovation of our approach lies in its iterative self-refining mechanism, which leverages spatial gene or protein expression patterns to progressively correct an initial coarse cellular map. Unlike conventional methods that treat segmentation as a one-off image processing task, SegJointGene learns from the underlying molecular biology of the tissue to define cell boundaries with better accuracy. A second key strength is its ability to perform joint segmentation and feature prioritization. In the process of delineating cells, SegJointGene simultaneously identifies a concise and biologically coherent set of genes or proteins that are most informative for defining cell types, providing valuable insights that are missed by other tools.

Despite its strong performance, SegJointGene has certain limitations that suggest avenues for future development. The iterative training process, while powerful, is computationally intensive and could be optimized for speed on large-scale datasets. Additionally, its performance is dependent on the quality of the initial nuclear segmentation, and errors in this first step could impact the final result.

SegJointGene can also be extended to analyze three-dimensional (3D) spatial datasets more efficiently, which may need new CNN architectures. Another direction is the integration of multi-modal data, where we could incorporate other spatial activities, such as chromatin accessibility (scATAC-seq) or epigenomic marks, to build a more comprehensive, multi-layered model of cellular and tissue organization.

## 5 Supplementary data

Supplementary data 1-3 are available as Excel files.

## 8 Data availability

The source code and SegJointGene framework are available at https://github.com/daifengwanglab/segjointgene. The mouse hippocampus dataset is available on Figshare at https://doi.org/10.6084/m9.figshare.7150760.

The Whole Mouse Brain dataset is hosted on the Brain Knowledge Platform at https://knowledge.brain-map.org/data/LVDBJAW8BI5YSS1QUBG. The human tonsil proteomics dataset can be accessed via Zenodo at https://doi.org/10.5281/zenodo.10982119.

## 9 Authors’ contributions

D.W. conceived the study. D.W. and H.M. designed the methodology and experiments. MH curated and processed the data required for the analysis. H.M. performed analysis and visualization. H.M. and D.W. wrote and edited the manuscript. All authors read and approved the final manuscript.

