## Supplementary Materials for "SegJointGene: joint cell segmentation and spatial gene prioritization by information entropy guided convolutional neural networks"

#### S1 Supplementary note 1: math variables

Table S1: Mathematical terms and concepts in SegJointGene

| **Variables** | **Definition** | **Description** |
|---|---|---|
| $\mathbf{A}$ | Initial segmentation map | A $w \times l$ matrix where $A_i \in \{1, \ldots, K\}$ denotes the cell-type label for pixel $i$. It serves as the current ground truth for the CNN in each iteration. |
| $\mathbf{A}'$ | Refined segmentation map | The updated segmentation map resulting from label re-assignment in the current iteration. It serves as the new initial segmentation map $\mathbf{A}$ for the next iteration. |
| $\mathbf{A}^*$ | Final segmentation map | The fully converged segmentation map returned as the final output of the algorithm after the iterative refinement process is complete. |
| $\mathbf{X}$ | Gene-density map stack | The 3D input tensor containing the density maps of $M$ genes. Derived from mRNA spot coordinates, this is the raw molecular feature input used by the CNN for segmentation learning. |
| $f(\cdot;\theta)$ | Convolutional neural network (CNN) | The function representation of the U-Net architecture, parameterized by weights $\theta$. It takes the gene-density maps $\mathbf{X}$ as input and outputs a $K$-dimensional probability map. |

| Variables | Definition | Description |
|---|---|---|
| $S_{jki}$ | **Pixel-specific Gene Importance Score for Cell-type Segmentation** | Defined as the negative entropy ($S_{jki} = -H_{jki}$), this metric quantifies the local relevance of a specific gene $j$ at spatial location (pixel) $i$ for the model's identification of cell type $k$. A higher score indicates that the expression of gene $j$ at pixel $i$ is critical for the prediction of cell type $k$, reflecting high model sensitivity to this specific feature. |
| $I_{ik}$ | **Aggregated Pixel Importance Score for Cell-type Segmentation** | The cumulative relevance score calculated by summing the pixel-specific scores across all genes $j$ for a given pixel $i$ and cell type $k$ ($I_{ik} = \sum_j S_{jki}$). This score represents the total evidence provided by the transcriptomic profile at pixel $i$ to support its assignment to cell type $k$, and is used as a thresholding criterion for boundary refinement. |
| $S_{jk}$ | **Gene Importance Score for Cell-type Segmentation** | The global importance metric for gene $j$ with respect to cell type $k$, derived by aggregating the pixel-specific scores over the entire spatial domain (all pixels $i$): $S_{jk} = \sum_i S_{jki}$. This final output captures the overall contribution of gene $j$ to defining cell type $k$ across the tissue, enabling the prioritization and ranking of spatially relevant genes. |

### S2 Supplementary note 2: Convocational neural network for segmentation

The **U-Net** is a convolutional neural network architecture originally developed for biomedical image segmentation, which has since become a de facto standard for a wide range of pixel-wise classification tasks ronneberger2015u. This design consists of a contracting path to capture context and a symmetric expanding path to enable precise localization, allowing the network to effectively process images and produce high-resolution segmentation masks.

#### S2.1 Encoder

The first part of the U-Net is the encoder, or contracting path, which follows the typical architecture of a convolutional network. Its primary purpose is to extract a hierarchical feature representation that captures the high-level **semantic context** of the image. This is achieved through a series of repeated blocks, each containing two 3x3 convolutions with ReLU activation functions, followed by a 2x2 max pooling operation for downsampling. With each downsampling step, the spatial dimensions of the feature maps are halved, while the number of feature channels is doubled. This process allows the network to build a rich, abstract feature representation and increase its receptive field to understand the image at different scales.

##### S2.2 Decoder

The second part of the architecture is the decoder, or expanding path. Its goal is to take the coarse, high-level feature map and progressively upsample it to reconstruct a full-resolution segmentation map, thereby recovering **precise spatial information** for localization. Each step in the decoder consists of an upsampling of the feature map using a transposed convolution, which doubles its spatial dimensions. This is immediately followed by a concatenation with the corresponding feature map from the encoder path via a skip connection, and then two standard 3x3 convolutions with ReLU.

##### S2.3 Skip connections

The most critical and innovative feature of the U-Net is its use of **skip connections**. These connections bridge the two paths by directly concatenating the feature maps from the contracting path to their corresponding layers in the expanding path. As the encoder downsamples, it loses fine-grained spatial information essential for creating precise boundaries in the final segmentation mask. The skip connections reintroduce this high-resolution information to the decoder. This fusion of shallow, high-resolution features from the contracting path with deep, semantic features from the expanding path allows the U-Net to make highly accurate predictions for each pixel, resulting in clean and well-defined object boundaries.

##### S2.4 U-Net architecture for segmentation

U-Net's architecture is exceptionally effective for segmentation because it elegantly solves the inherent tension in the task: the need to both understand the broader context of the image and make a precise decision for every single pixel. The encoder path provides the necessary contextual understanding, while the decoder path with its skip connections ensures that the fine-grained spatial detail needed for accurate localization is not lost. This dual capability makes it a powerful and versatile tool, leading to its widespread adoption in scientific and industrial image analysis.

#### S3 Supplementary figures

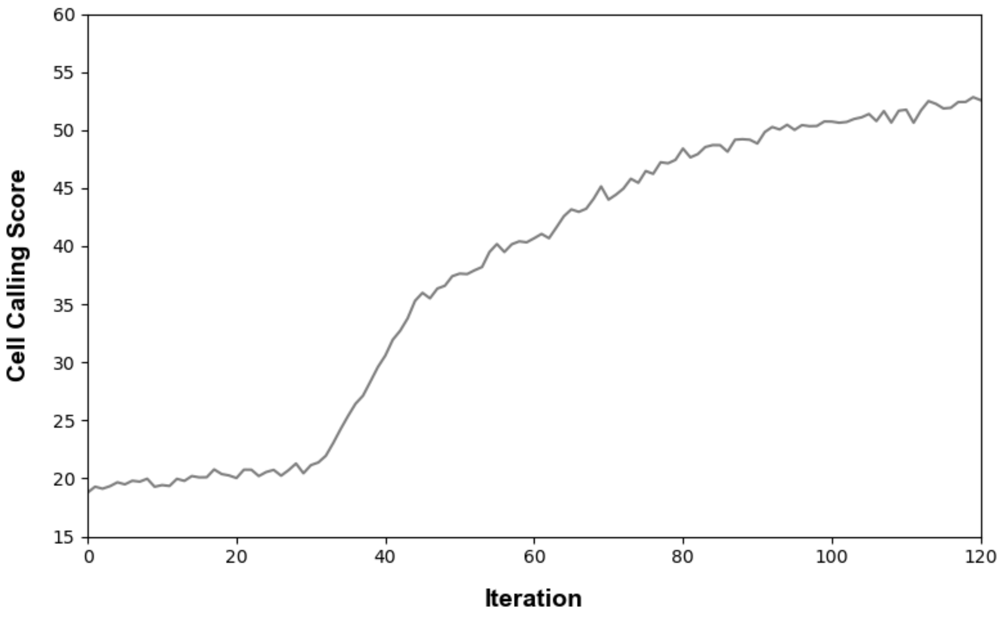


Figure S1: **SegJointGene training aims to increase cell calling score and finally converge through iterations, such as for the mouse hippocampus data.**

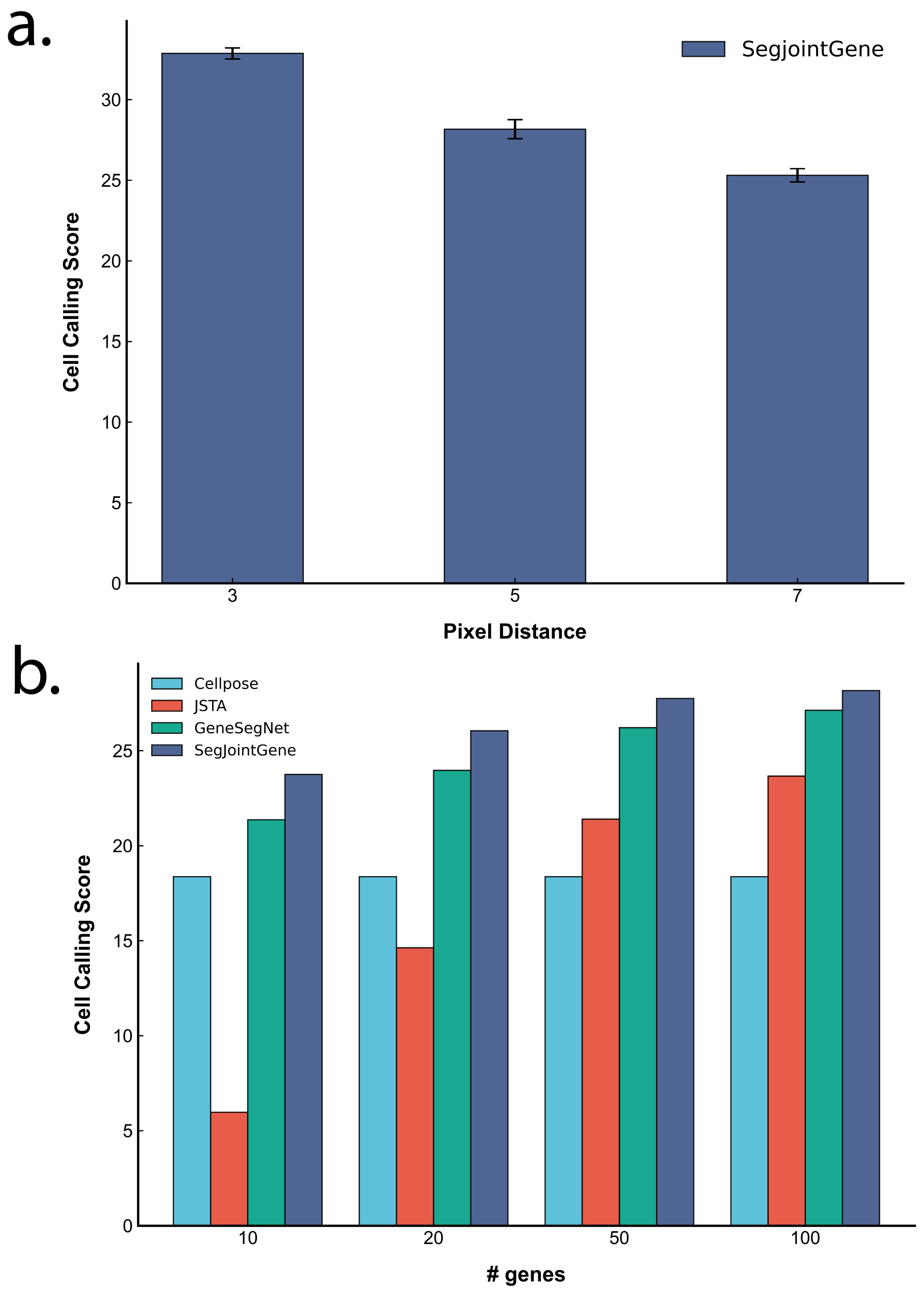


Figure S2: **Robustness evaluation of SegJointGene with varying numbers of input genes.** (**a**) Cell calling score of SegJointGene across different pixel distances, with error bars representing the standard deviation from five independent runs. (**b**) Quantitative comparison of SegJointGene against existing segmentation approaches using the cell calling score metric for the mouse prefrontal cortex data. The evaluation was performed at a fixed pixel-distance threshold of 5 pixels, while varying the number of input genes used for the model.

#### S4 Supplementary data 1

**Description:** The Excel file contains the SegJointGene gene importance scores for the mouse hippocampus dataset (**Fig. 3**).

#### S5 Supplementary data 2

**Description:** The Excel file provides the SegJointGene gene importance scores for the mouse prefront cortex from the Whole Mouse Brain dataset (**Fig. 4**).

#### S6 Supplementary data 3

**Description:** The Excel file contains the SegJointGene protein importance scores for the human tonsil dataset (**Fig. 5**).
